# Evaluation of maternal-infant dyad inflammatory cytokines in pregnancies affected by maternal SARS-CoV-2 infection in early and late gestation

**DOI:** 10.1101/2021.12.26.472655

**Authors:** Elizabeth S. Taglauer, Yashoda Dhole, Jeffery Boateng, Jennifer Snyder-Cappione, Samantha E. Parker, Katherine Clarke, Lillian Juttukonda, Jean Devera, Jessica Hunnewell, Elizabeth Barnett, Hongpeng Jia, Christina Yarrington, Vishakha Sabharwal, Elisha M. Wachman

## Abstract

**Objective:** SARS-CoV-2 infection induces significant inflammatory cytokine production in adults, but infant cytokine signatures in pregnancies affected by maternal SARS-CoV-2 are less well characterized. We aimed to evaluate cytokine profiles of mothers and their infants following COVID-19 in pregnancy.

**Study Design:** Serum samples at delivery from 31 mother-infant dyads with maternal SARS-CoV-2 infection in pregnancy (COVID) were examined in comparison to 29 control dyads (Control). Samples were evaluated using a 13-plex cytokine assay.

**Results:** In comparison with controls, interleukin (IL)-6 and interferon gamma-induced protein 10 (IP-10) were higher in COVID maternal and infant samples (p<0.05) and IL-8 uniquely elevated in COVID infant samples (p<0.05). Significant elevations in IL-6, IP-10 and IL-8 were found among both early (1^st^/2^nd^ Trimester) and late (3^rd^ Trimester) maternal SARS-CoV-2 infections.

**Conclusions:** Maternal SARS-CoV-2 infections throughout gestation are associated with increased maternal and infant inflammatory cytokines at birth with potential to impact long-term infant health.

## INTRODUCTION

COVID-19 has impacted a growing number of pregnant patients throughout the pandemic, affecting an estimated 14 per 1000 births in the U.S. in 2020, with yet undefined risks for long-term infant health (1). Throughout the pandemic, clinical studies have consistently shown that maternal SARS-CoV-2 infection during pregnancy generally does not result in vertical transmission from mother to infant (2–4). While the rarity of vertical SARS-CoV-2 transmission has been well-characterized, the effects of maternal SARS-CoV-2 infection in pregnancy on the developing fetus are less well defined. Of particular importance is the maternal immune activation and inflammation in response to SARS-CoV-2 which has significant potential to cause harm to the developing fetus. Indeed, inflammatory exposures during in the perinatal period have been shown to impair infant growth and development in other disease processes (5–7) with a particular risk for alterations in brain development and cognitive function later in life (8).

In acute COVID-19 infections, intense immune activation leads to an overproduction of serum cytokines known as a “cytokine storm”, with significant increases in multiple inflammatory mediators including IL-1, IL-6, tumor necrosis factor (TNF)-α, interferon (IFN)-γ and IP-10 (9, 10). While pregnant COVID-19 patients have a variety of clinical presentations, SARS-CoV-2 infection in pregnancy has been associated with increased COVID-19 disease severity and higher rates of intensive care unit (ICU) admissions when compared with COVID-19 in non-pregnant patients. Further, in comparison to pregnant patients without SARS-CoV-2 infections, pregnant patients with COVID-19 have an increased risk for preeclampsia and increased maternal mortality (11, 12). Additionally, there is substantial evidence supporting intrauterine inflammation in pregnancies affected by maternal COVID-19 disease. Multiple studies have identified histological evidence of placental inflammatory lesions (13, 14) and male fetus-specific upregulation of placental interferon response genes and pro-inflammatory cytokines/chemokines (15).

More recent publications have also identified altered expression of maternal inflammatory cytokine profiles and evidence of fetal leukocyte activation through cord blood analysis in limited sample cohorts from pregnancies with COVID-19 in the third trimester (16, 17). However, no study to date has characterized corresponding maternal and infant cytokine repertoires in larger cohorts, particularly from pregnancies with maternal SARS-CoV-2 infections in early pregnancy. Here we hypothesized that when compared to control pregnancies, SARS-CoV-2 positive mothers and their infants have significant alterations in inflammatory cytokines and that these alterations vary by the gestational timing of maternal COVID-19. In the current study, serum cytokines were evaluated in mother-infant dyads by examining maternal blood and cord blood / infant blood at the time of delivery between SARS-CoV-2 positive and control pregnancies (i.e. contemporary patients negative for SARS-CoV-2 at delivery). In particular, we evaluated changes relative to the gestational timing of maternal SARS-CoV-2 infection.

## SUBJECTS AND METHODS

### Study Design and Patient Population

Boston Medical Center (BMC) is the largest urban safety-net hospital in New England serving most of the underrepresented minority populations living in the Greater Boston Area. This hospital delivers a significant proportion of pregnant patients from minority groups with comorbidities such as hypertension and obesity and having public insurance, which are characteristics associated with severe SARS-CoV-2 disease (18, 19). BMC performed universal nasopharyngeal swab PCR screening of all pregnant patients who presented in labor starting in April 2020.

We enrolled mother-infant dyads from the obstetric prenatal clinics, Labor and Delivery, and the Postpartum Unit in a prospective cohort study from July 2020 through June 2021. To be eligible for the COVID-19 group, patients had to be at least 18 years of age, have a positive SARS-CoV-2 infection at any point during pregnancy as documented by nasopharyngeal swab PCR testing, have a viable singleton gestation pregnancy, and be English or Spanish speaking. Patients were excluded from this cohort if they had received COVID-19 vaccination by the time of delivery, or if they were deemed unable to provide informed consent and agree to study procedures for any medical or social reason.

The control group was enrolled between January and April 2021. Eligibility criteria for the control group were age at least 18 years of age, documented negative SARS-CoV-2 PCR testing throughout their pregnancy without suspected COVID-19, a viable singleton gestation pregnancy, and English or Spanish speaking. Of note, this group was also unvaccinated for SARS-CoV-2.

This study was approved by the Boston University Medical Campus Institutional Review Board. Written informed consent or REDCap e-consent was obtained from all participants. Institutional COVID-19 research protocols were followed with use of a remote consent process and coordination with clinical care blood draws for patients on COVID-19 precautions at the time of delivery.

### Sample Collection

Maternal blood samples (3mL in an EDTA tube) were collected by L&D staff or phlebotomy trained research staff. Cord blood samples (3-5mL in an EDTA tube) were collected by L&D staff in the delivery room. If cord blood could not be obtained, an infant blood sample (0.5-1mL in a EDTA microtainer) was obtained via heel stick by trained neonatal study staff or bedside nursing staff. Blood was centrifuged within 6 hours of collection in our BSL-2+ research laboratory space using COVID-19 specific safety protocols. Plasma was then extracted, aliquoted into 0.5mL aliquots, and frozen at −80°C until analysis.

### Cytokine analysis

Serum cytokine levels were evaluated using a flow cytometry bead-based LEGENDplex assay (BioLegend). Frozen plasma samples were quick-thawed and analyzed using a13-plex human anti-viral response panel to analyze the following cytokines: IP-10, IL-1β, IL-6, TNF-α, IFN-λ1, IL-8, IL-12p70, IFN-α2, IFN-λ 2/3, GM-CSF, IFN-β, IL-10 and IFN-γ. Samples were pre-diluted 1:2 and the kit was run according to manufacturer’s instructions. Sample data were collected using the Cytek^®^ Aurora flow cytometer (Cytek Biosciences) in the Boston University School of Medicine Flow Cytometry Core facility. Cytokine concentrations for each sample were determined from extrapolation from standard curves using the LEGENDplex Data Analysis Software. Pilot testing of 20 cord blood and 20 infant blood samples indicated a very high correlation for this cytokine panel, thus these sample types were grouped together for the purposes of analysis as an “infant” sample for analysis of the full cohort.

### Data Abstraction

The electronic medical records of the mother-infant dyads enrolled were reviewed. Demographic characteristics such as race, ethnicity, age at delivery, and primary language were recorded in a secure electronic study database. Other maternal data points such as history of chronic illnesses, pregnancy related diagnosis and SARS-CoV-2 infection related symptoms were abstracted. Trimester and gestational age at the time of infection, severity of infection (hospitalization / ICU), and maternal COVID-19 status at the time of delivery were also collected. For infants, demographics and birth parameters as well as COVID-19 testing status and neonatal symptoms in the first 30 days of life were reviewed. Infants were also tracked for any emergency room visits or readmissions in the first 30 days of life.

### Statistical Methods

Demographics between the COVID-19 and control groups were compared using t-tests and chi-square as indicated. For cytokine statistical analysis, due to a large proportion of non-detects in several of the cytokines examined, a Kaplan Meier method for left-censored data was used for cytokine statistical analysis. This is a non-parametric technique that estimates a probability distribution based on the number of samples below detected concentrations. (20). This method is preferred over simple substitution methods particularly when there is a large proportion of censored data. This Kaplan Meier method was used to calculate means and standard deviations separately within COVID and control groups and by sample type (maternal blood and cord /infant blood). Mean differences and 95% confidence intervals between the COVID and reference groups were then calculated for each cytokine within both maternal and cord/infant samples. We also examined differences between mean cytokine levels in maternal blood and cord blood / infant blood by timing of SARS-CoV-2 infection during pregnancy. Mean cytokine levels for early COVID (1^st^/2^nd^ trimester) and late COVID (3^rd^ trimester) were compared to each other and then to the reference group using t-tests for independent samples with unequal variances. Lastly, among the COVID-19 dyads, we calculated the correlation coefficient for individual cytokines between maternal and cord/infant samples. All statistical analysis was performed using SAS software V9.4 (SAS Analytics). Graphical data for cytokine levels was created using Prism Software (Graphpad)

## RESULTS

### Cohort Clinical Data

For our prospective study design (**Figure 1**) we enrolled pregnant patients with SARS-CoV-2 in early (1^st^/2^nd^ trimester) and late (3^rd^ trimester) gestational stages of pregnancy along with contemporary controls who had no documented history of COVID-19 and also negative SARS-CoV-2 testing at time of delivery. We identified 146 pregnant individuals with COVID-19 who delivered infants at BMC between July 2020 and April 2021. Of those, 20 were not eligible for approach for the study due to twin gestation (n=1), non-English or Spanish speaking (n=11), severe social concerns screened out by providers (n=3), maternal age < 18 years (n=3), or ICU admission during screening (n=2). Of the remaining 126 patients, 85 were approached for consent. The remainder were not approached due to availability of research staff and timing during labor (no patient in advanced stages of labor or immediately after delivery were approached). Of those 85 patients, 60 mothers (71%) provided informed consent for the study. From those 60 consented, 31 dyads with complete maternal blood and cord blood matched samples from delivery were selected at random for the cytokine analysis.

**Figure 1.**
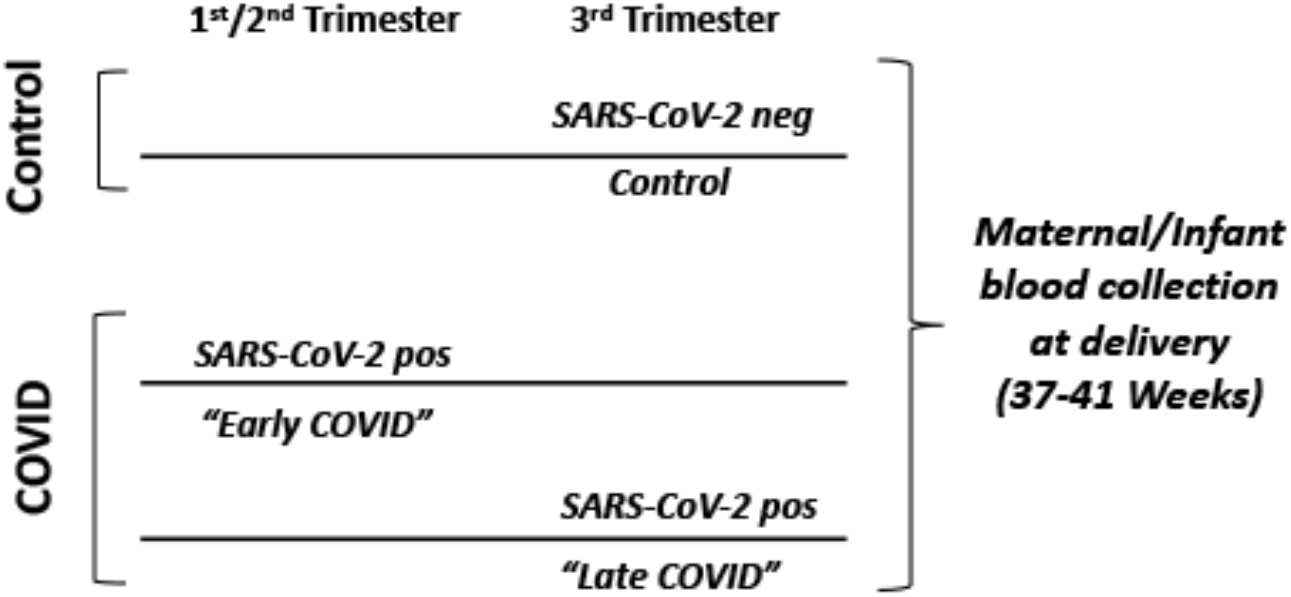
Study Design. Study design of patient cohorts for serum cytokine analysis. *Control*: mothers with no report of COVID-19 disease at any time during their pregnancy and negative for SARS-CoV-2 at time of delivery screening. *Early COVID*: mothers with SARS-CoV-2 positive testing within the 1^st^ or 2^nd^ trimester of pregnancy (1-27 weeks gestation). *Late COVID*: mothers with SARS-CoV-2 positive testing in the 3^rd^ trimester of pregnancy (28-41weeks gestation).

For our contemporary control cohort recruitment during the same time period, a total of 256 patients were admitted in labor. Of those, 20 were identified as COVID-19 positive thus ineligible, and others screened out due to non-English or Spanish speaking (n=23), being screened out by providers due to social concerns (n=5), or other medical reasons for ineligibility such as twin gestation (n=6). Of the remaining 202 patients, 41 (20%) were approached with our goal of enrolling 2 control dyads per week due to staff availability and timing of labor. Of the 41 patients approached, 31 (75%) consented for the study. Of those 31 dyads, 29 had complete maternal blood and cord blood matched samples thus were included in the analysis.

Demographics of the COVID and control cohorts are shown in **Table 1**. There were no significant differences in maternal age, race, ethnicity, language, delivery mode, gestational age, birth weight, infant sex, or infant outcomes between the two cohorts. The COVID-19 cohort contained 3 patients (9.7%) with infections in the first trimester, 18 patients (58.1%) with infections in the 2^nd^ trimester and 10 patients (32.3%) with infections in the third trimester. Twenty-eight patients (90.3%) in our COVID cohort had documentation of COVID-19 symptoms at any stage during their pregnancy with 6.6% requiring hospitalization for COVID-19 during pregnancy. Some differences were noted in the presence of various pregnancy co-morbidities and chronic health conditions, with the most significant being intrauterine growth restriction in the COVID-19 group. None of the infants were diagnosed with SARS-CoV-2 infection within 30 days of delivery.

**Table 1:** Demographics of COVID-19 versus control mother-infant dyads.

| Variable | Mean (SD) or N(%)<br>CONTROL<br>N=28 | Mean (SD) or N(%)<br>COVID-19<br>N=31 | P-Value |
| --- | --- | --- | --- |
| Maternal age at delivery (years) | 29.8 (5.7) | 29.7 (5.6) | 0.96 |
| Maternal Race |  |  | 0.17 |
| Black | 5 (17.9%) | 2 (6.5%) |  |
| White | 5 (17.9%) | 8 (25.8%) |  |
| Asian | 1 (3.6%) | 2 (6.5%) |  |
| Other | 14 (50.0%) | 19 (61.3%) |  |
| Missing | 3 (10.7%) | 0 (0%) |  |
| Maternal Ethnicity |  |  | 0.41 |
| Hispanic | 18 (64.3%) | 23 (74.2%) |  |
| Non-Hispanic | 10 (35.7%) | 8 (25.8%) |  |
| Unknown | 0 (0%) | 0 (0%) |  |
| Maternal Primary Language |  |  | 0.14 |
| English | 18 (64.3%) | 14 (45.2%) |  |
| Spanish | 10 (35.7%) | 17 (54.8%) |  |
| Health Insurance |  |  | 0.60 |
| Public | 20 (71.4%) | 24 (77.4%) |  |
| Private | 8 (28.6%) | 7 (22.6%) |  |
| Chronic health conditions |  |  |  |
| Any condition | 17 (60.7%) | 21 (67.7%) | 0.57 |
| Diabetes | 0 (0%) | 0 (0%) | N/A |
| Hepatitis C | 2 (7.1%) | 0 (0%) | 0.13 |
| Hypertension | 2 (7.1%) | 1 (3.2%) | 0.49 |
| Obesity | 4 (14.3%) | 3 (9.7%) | 0.58 |
| Thyroid condition | 4 (14.3%) | 1 (3.2%) | 0.13 |
| Substance use disorder | 2 (7.1%) | 1 (3.2%) | 0.49 |
| Other | 15 (53.6%) | 13 (41.9%) | 0.37 |
| Pregnancy co-morbidities |  |  |  |
| Any co-morbidity | 23 (82.1%) | 29 (93.6%) | 0.17 |
| Chorioamnionitis | 0 (0%) | 4 (12.9%) | 0.05 |
| Gestational diabetes | 6 (21.4%) | 4 (12.9%) | 0.38 |
| Preeclampsia/Gestational hypertension | 10 (35.7%) | 9 (29.0%) | 0.58 |
| Intrauterine growth restriction | 1 (3.6%) | 7 (22.6%) | **0.03** |
| Preterm labor | 0 (0%) | 2 (6.5%) | 0.17 |
| Other | 16 (57.1%) | 21 (67.7%) | 0.40 |
| Nicotine smoking | 2 (7.1%) | 2 (6.5%) | 0.31 |
| Delivery mode |  |  |  |
| Vaginal | 22 (78.6%) | 22 (71.0%) | 0.50 |
| C-section | 6 (21.4%) | 9 (29.0%) |  |
| Gestational age at delivery (weeks) | 39.1 (1.3) | 39.3 (1.6) | 0.50 |
| Infant birth weight (grams) | 3349 (513.9) | 3306 (547.4) | 0.76 |
| Infant sex |  |  |  |
| Male | 15 (53.6%) | 16 (51.6%) | 0.69 |
| Female | 13 (46.4%) | 15 (48.4%) |  |
| Breastfed infant | 26 (92.9%) | 30 (96.8%) | 0.49 |
| 5 minute APGAR score | 8.8 (0.6) | 8.9 (0.5) | 0.32 |
| Infant length of hospitalization (days) | 3.5 (3.0) | 3.1 (2.1) | 0.59 |
| Infant ER visit within 30 days | 3 (11.1%) | 1 (3.2%) | 0.24 |
| Infant re-hospitalization with 30 days | 3 (11.1%) | 1 (3.2%) | 0.24 |
| Infant diagnosed with SARS-CoV-2 within 30 days of delivery | 0 (0%) | 0 (0%) | N/A |
| Mother diagnosed with SARS-CoV-2 within 30 days of delivery | 0 (0%) | N/A | N/A |
| Trimester of COVID-19 infection<br>First<br>Second<br>Third | N/A | 3 (9.7%)<br>18 (58.1%)<br>10 (32.3%) | N/A |
| Gestational age at infection (weeks) | N/A | 24.0 (9.0) | N/A |
| *Maternal symptoms of COVID-19 at time of SARS-CoV-2 positive testing*<br>No<br>Yes | N/A | 3 (9.7%)<br>28 (90.3%) | N/A |
| *Maternal COVID-19 symptoms at delivery*<br>Confirmed SARS-CoV-2 with active symptoms at delivery<br>Confirmed SARS-CoV-2 and asymptomatic at delivery<br>Recovered from SARS-CoV-2 infection earlier in pregnancy (1<sup>st</sup>, 2<sup>nd</sup> and 3<sup>rd</sup> Trimesters) | N/A | 1 (3.2%)<br>4 (12.9%)<br>26 (83.9%) | N/A |
| Hospitalized for COVID-19<br>No<br>Yes | N/A | 29 (93.6%)<br>2 (6.6%) | N/A |

**Abbreviations:** ER = emergency room; SARS-CoV-2 = Severe acute respiratory syndrome coronavirus 2

### Serum cytokine analysis

We used a multi-plex approach to plasma analysis. A viral response cytokine panel was selected which included cytokines previously identified to be altered among adult COVID-19 patients (9, 10) as well as cytokines associated with adverse long term fetal outcomes in perinatal inflammatory exposures (5–8). These included IP-10, IL-1β, IL-6, TNF-α, IFN-λ1, IL-8, IL-12p70, IFN-α2, IFN-λ 2/3, GM-CSF, IFN-β, IL-10 and IFN-γ. Maternal and infant cytokine levels were compared between control and COVID-19 cohorts (**Figures 2–4, Supplemental Table 1**) with additional sub-analysis of Control (SARS-CoV-2 negative) vs Early COVID (1^st^/2^nd^ Trimester) and Late COVID (3^rd^ Trimester) maternal SARS-CoV-2 infections in pregnancy (**Figures 2–4, Supplemental Table 2**). Finally, we conducted a correlation analysis of maternal-infant cytokine levels in maternal-infant dyads within the COVID-19 cohort (**Figures 2–4, Supplemental Table 3**).

**Figure 2:**
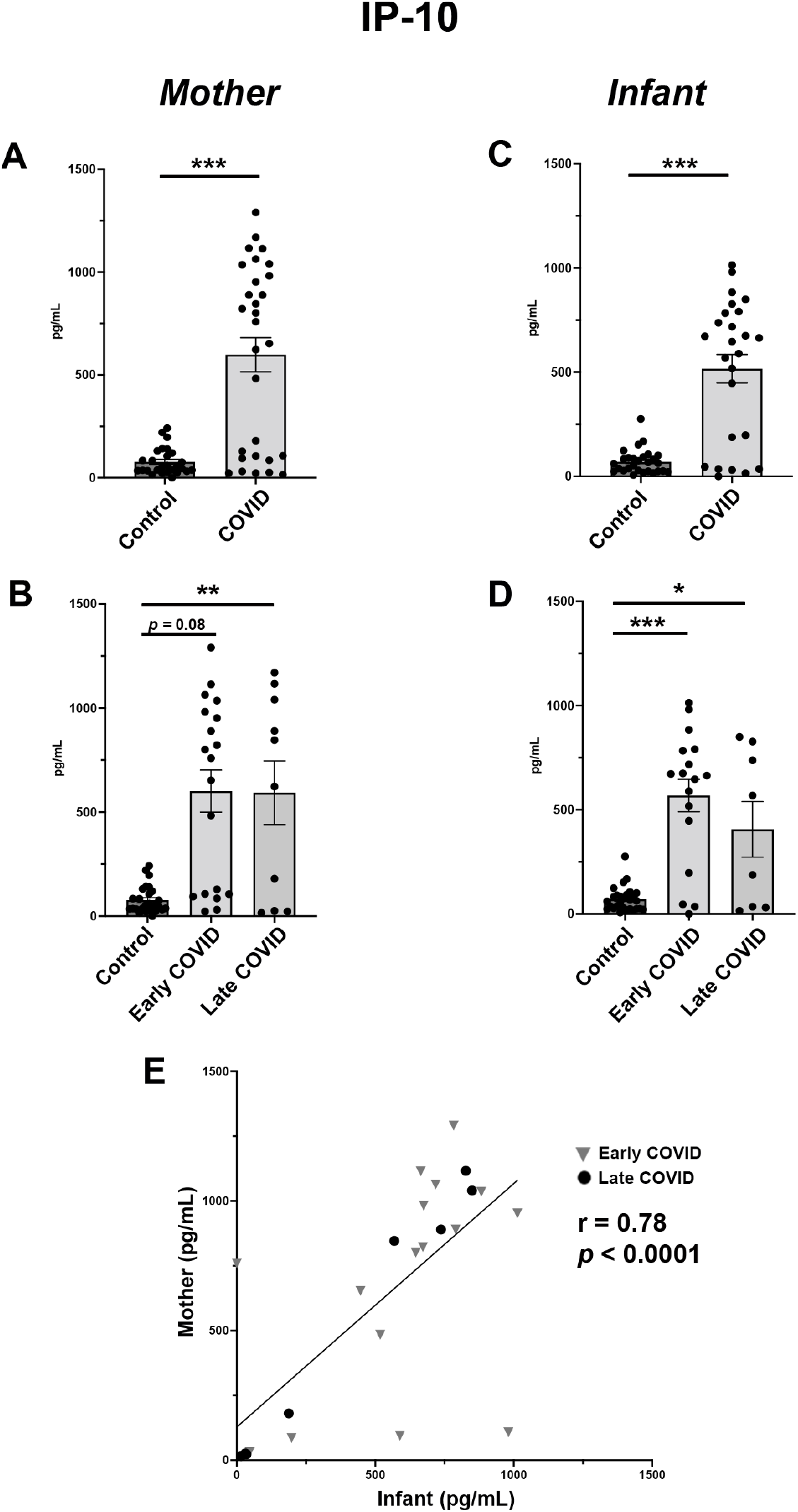
IP-10 is significantly elevated in maternal and infant serum following COVID-19 infections in pregnancy. A: Maternal serum IP-10 expression in Control vs COVID cohorts. B: Maternal serum IP-10 sub-analysis among Control, Early COVID and Late COVID cohorts. C. Infant serum IP-10 expression in Control vs COVID cohorts. D: Infant serum IP-10 sub-analysis among Control, Early COVID and Late COVID cohorts. E: Correlation analysis of IP-10 expression in maternal-infant dyads. *Control*: as described in Figure 1. *COVID*: Mothers positive for SARS-CoV-2 in pregnancy. *Early COVID*: as described in Figure 1. *Late COVID*: as described in Figure 1. * *p* < 0.05, ** *p* < 0.01, \*\*\**p* <0.001

**Figure 3:**
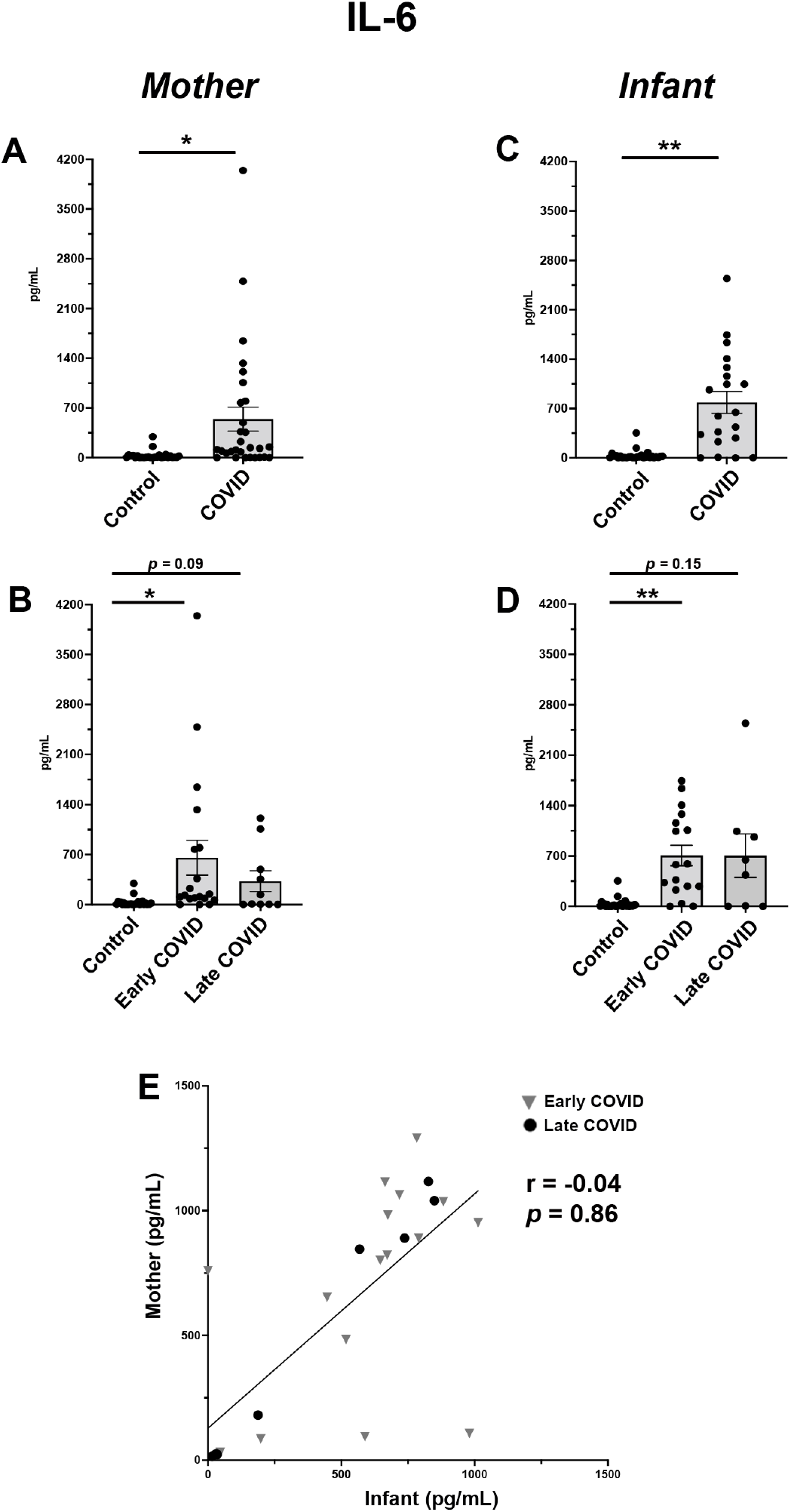
Maternal and infant serum IL-6 levels are distinctly elevated in SARS-CoV-2 infections early in pregnancy. A: Maternal serum IL-6 expression in Control vs COVID cohorts. B: Maternal serum IL-6 sub-analysis among Control, Early COVID and Late COVID cohorts. C. Infant serum IL-6 expression in Control vs COVID cohorts. D: Infant serum IL-6 sub-analysis among Control, Early COVID and Late COVID cohorts. E: Correlation analysis of IL-6 expression in maternal-infant dyads. *Control*, *COVID*, *Early COVID*, and *Late COVID*: as described in Figures 1 and 2. * *p* < 0.05, ** *p* < 0.01

**Figure 4:**
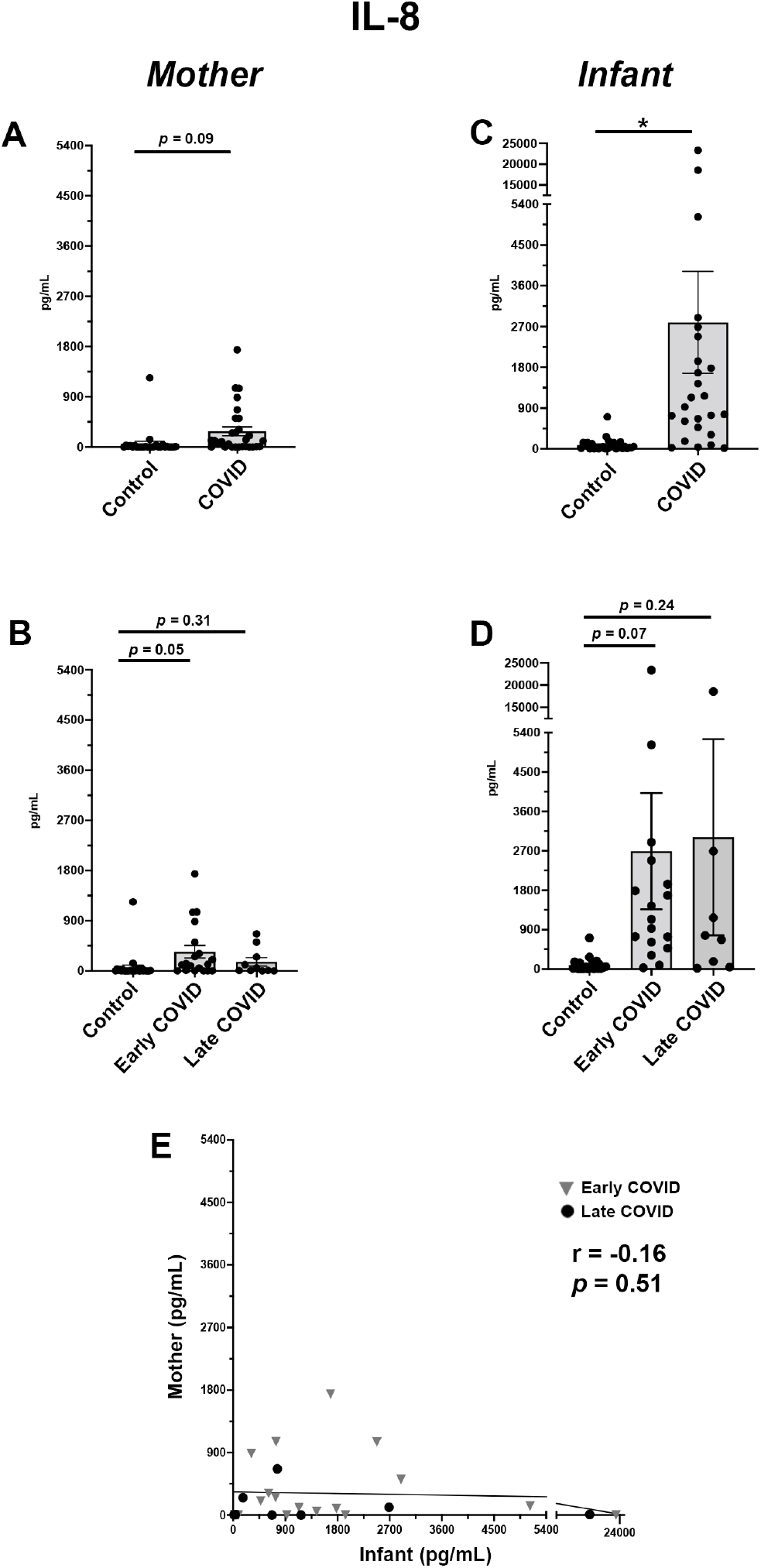
Uniquely elevated levels of IL-8 in serum of infants from pregnancies affected by maternal SARS-CoV-2 in early pregnancy. A: Maternal serum IL-8 expression in Control vs COVID cohorts. B: Maternal serum IL-8 sub-analysis among Control, Early COVID and Late COVID cohorts. C. Infant serum IL-8 expression in Control vs COVID cohorts. D: Infant serum IL-8 sub-analysis among Control, Early COVID and Late COVID cohorts. E: Correlation analysis of IL-8 expression in maternal-infant dyads. *Control*, *COVID*, *Early COVID*, and *Late COVID*: as described in Figures 1 and 2. * *p* < 0.05

Among all analytes, those with the most significant changes were pro-inflammatory cytokines IP-10, IL-6 and IL-8. For IP-10, both maternal and infant blood showed significantly higher levels in the COVID cohort as compared with Controls (**Figure 2 A,C**). Interestingly, these elevations were noted in both Early COVID and Late COVID groups. While the most significant elevations in IP-10 maternal levels were found to be in the Late COVID group (**Figure 2B**), the highest infant blood levels were noted in Early COVID pregnancies (**Figure 2D**). Regression analysis further demonstrated a positive correlation in maternal and infant IP-10 levels among dyads analyzed (**Figure 2E**).

IL-6 analysis also showed significant upregulation in both maternal and infant blood from COVID pregnancies compared with Controls (**Figure 3 A,C**). When analyzing by gestational stage of SARS-CoV-2 infection, we identified that the Early COVID cohort had the highest IL-6 elevations for both maternal and infant samples (**Figure 3 B,D**). However, these elevations in IL-6 were not significantly correlated within individual mother-infant dyads (**Figure 3E**).

When evaluating IL-8, only infant samples showed a significant elevation in the COVID cohort as compared with controls (**Figure 4C**). While this was significant for total cohort analysis, these values did not reach statistical significance in sub-analysis by gestational stage of SARS-CoV-2 infection (**Figure 4D**), likely due to the significant variability in cytokine values among Early and Late COVID sub-groups. Further, the abundance of IL-8 was also notably higher in the infant COVID cohort (**Figure 4C**) as compared with the maternal COVID cohort (**Figure 4A**). As expected, there was no correlation between maternal and infant cytokine levels among dyads (**Figure 4E).**

Among the remaining cytokines, IFN-λ1 showed a trend in elevation among COVID maternal samples, particularly among the Early COVID cohort (**Supplemental Figure 1 A,B**). While regression analysis did show significant correlation among dyads (**Supplemental Figure 1E)**, the correlation significance appeared to originate in highly elevated levels from two dyads with the remainder of the dyads having minimal cytokine values among the infant samples. All other cytokines assayed showed no significant differences in COVID vs Control group comparisons among maternal and infant samples (**Supplemental Table 1, 2 and 3**).

## DISCUSSION

This study identified a unique repertoire of inflammatory cytokines with significant alterations in maternal-infant dyads from pregnancies affected by maternal SARS-CoV-2 infection, namely IP-10, IL-6 and IL-8. Importantly, these alterations were identified even among the Early COVID cohort, suggesting persistent elevations of inflammatory cytokines in maternal and neonatal circulation months after initial maternal SARS-CoV-2 infection.

The COVID-associated maternal serum elevations of IP-10 and IL-6 were expected findings as both are central components of the COVID-19 cytokine storm (9, 10, 21) and 90% of our maternal COVID-19 cohort was symptomatic at the time of the positive SARS-CoV-2 tests, indicating the majority of our cohort had active disease rather than asymptomatic carrier status. These data also suggest a prolonged upregulation of these cytokines lasting weeks to months after disease onset as as only a small percentage (1%) had active symptoms at the time of delivery sample collection. The differing maternal-infant correlation profiles for IP-10, IL-6 and IL-8 between mothers and infants in the COVID cohort suggest that the infant cytokine elevations are not merely a passive transfer of maternal cytokines passing into fetal circulation. Rather, these changes may be the result of an independent fetal immune response to maternal SARS-CoV-2 in pregnancy, as also suggested by a recent study showing evidence of fetal leukocyte activation in a cohort (n=3) of cord blood samples from pregnancies affected by maternal COVID-19 in the third trimester (17).

IP-10 (also known as CXCL10) is an established inflammatory chemokine secreted by both immune and parenchymal cell types with varied functions to induce leukocyte chemotaxis, trigger cellular apoptosis, and inhibit vascular growth (22). In adult COVID-19 patients, IP-10 has a long-lasting serum elevation profile that is unique from its secretion pattern in other viral infections (21). Our data were congruent with these findings showing persistent IP-10 elevations in maternal and most significantly infant serum from early gestational COVID-19 infections. In pregnancy, elevated maternal IP-10 has been implicated in miscarriage and preeclampsia (23, 24), but the long-term infant effects of IP-10 exposure in the perinatal period are currently undefined.

IL-6 and IL-8 are also both well-characterized inflammatory mediators central in the COVID-19 cytokine response (9, 10). In pregnancy, elevated serum levels of IL-6 and IL-8 have been associated with gestational pathologies including miscarriage, preeclampsia and preterm delivery (25, 26). Perinatal exposure to these cytokines has been associated with altered fetal development. In preclinical studies, IL-6 is a central mediator of enhanced postnatal offspring intestinal inflammatory T cell activation following maternal perinatal infection (27). Additionally, IL-6 and IL-8 elevations in pregnancy have independently been associated with altered infant neurodevelopmental outcomes (28, 29), with prenatal exposures to IL-8 particularly implicated in the risk of schizophrenia later in life (30, 31).

The current study has several limitations. First, our cohorts were of moderate sample size and we did not evaluate a complete repertoire of COVID-19 related clinical variables (i.e. c-reactive protein (CRP) levels, white blood cell count). As only a small subset of our cohort had these labs available, they were not included as part of this analysis. Our cohort also had a larger proportion of patients with maternal SARS-CoV-2 infections in early gestation as compared with late gestation. Ongoing analysis with larger cohorts will be required to characterize the cytokine profiles of early vs late pregnancy SARS-CoV-2 infections more completely. As the majority of studies on COVID-19 in pregnancy contain samples exclusively from late pregnancy (3^rd^ trimester) infections, our study highlights the importance of incorporating patients with SARS-CoV-2 infection at multiple gestational stages of pregnancy for ongoing studies in this field. In particular, multivariate analysis to correlate anti-viral antibodies with cytokine profiles will be informative to identify how these inflammatory signatures impact maternal and infant SARS-CoV-2 responses, particularly in relation to gestational timing of maternal infection.

This study supports a growing body of evidence that perinatal alterations resulting from maternal COVID-19 in pregnancy have a risk of impacting the health of infants even in the absence of fetal SARS-CoV-2 transmission. Indeed, of all our clinical parameters evaluated between control and COVID cohorts, the only value which reached statistical significance was an increase in fetal intrauterine growth restriction (IUGR) in the COVID cohort. As there was no accompanying difference in the incidence of preeclampsia, the growth restriction could be an early indicator of primary fetal pathology resulting from altered intrauterine physiology in response to maternal SARS-CoV-2 infection. Future studies involving more extensive patient cohorts and pre-clinical models will be required to characterize the driving mechanisms and developmental impact of the cytokine alterations identified in this study. Finally, our work highlights the importance of long-term follow-up for infants from pregnancies affected by maternal SARS-CoV-2 as an at-risk population of the COVID-19 era.

## ACKNOWLEDGMENTS

We would like to acknowledge all of the bedside nurses, midwives, and physicians from the Boston Medical Center Labor and Delivery Unit, Postpartum Unit, and Newborn Intensive Care Unit who assisted us with subject recruitment and facilitation of sample collection, especially Kate Thibault, RN; Lauren Laliberte, RN; Brianna Medeiros, RN, NP; Elizabeth Regan, RN, and Joanna Bushfield, RN; Sigride Jean-Sicard, RN, NP; Elizabeth Woodard, RN, NP; Alice Cruikshank, RN, NP; Bharati Sinha, MD; Ruby Bartolome, DO; and Margaret G. Parker, MD, MPH. We would also like to acknowledge the Maxwell Finland Laboratory of Pediatric Infectious Disease at Boston Medical Center, and the Boston University laboratory of Dr. Jennifer Snyder-Cappione.

## AUTHOR CONTRIBUTIONS

EW, EB, CY, VS, and ET were involved in development of the study protocol and procedures. JB, LJ, ET, and EW enrolled subjects in the study. YD, JB, LJ, ET, and EW collected and processed samples for the study. KC and JC performed the cytokine assays for the study. YD and JD performed the chart abstraction for the study. JH and SP performed the statistical data analysis for the study. ET, YD and EW were involved in data interpretation, figure composition and final manuscript composition. EW provided overall oversight and obtained grant funding for the study. All authors reviewed and edited the manuscript and provided final approval of the submitted version.

## SUPPLEMENTAL DATA

**Supplemental Figure 1.**
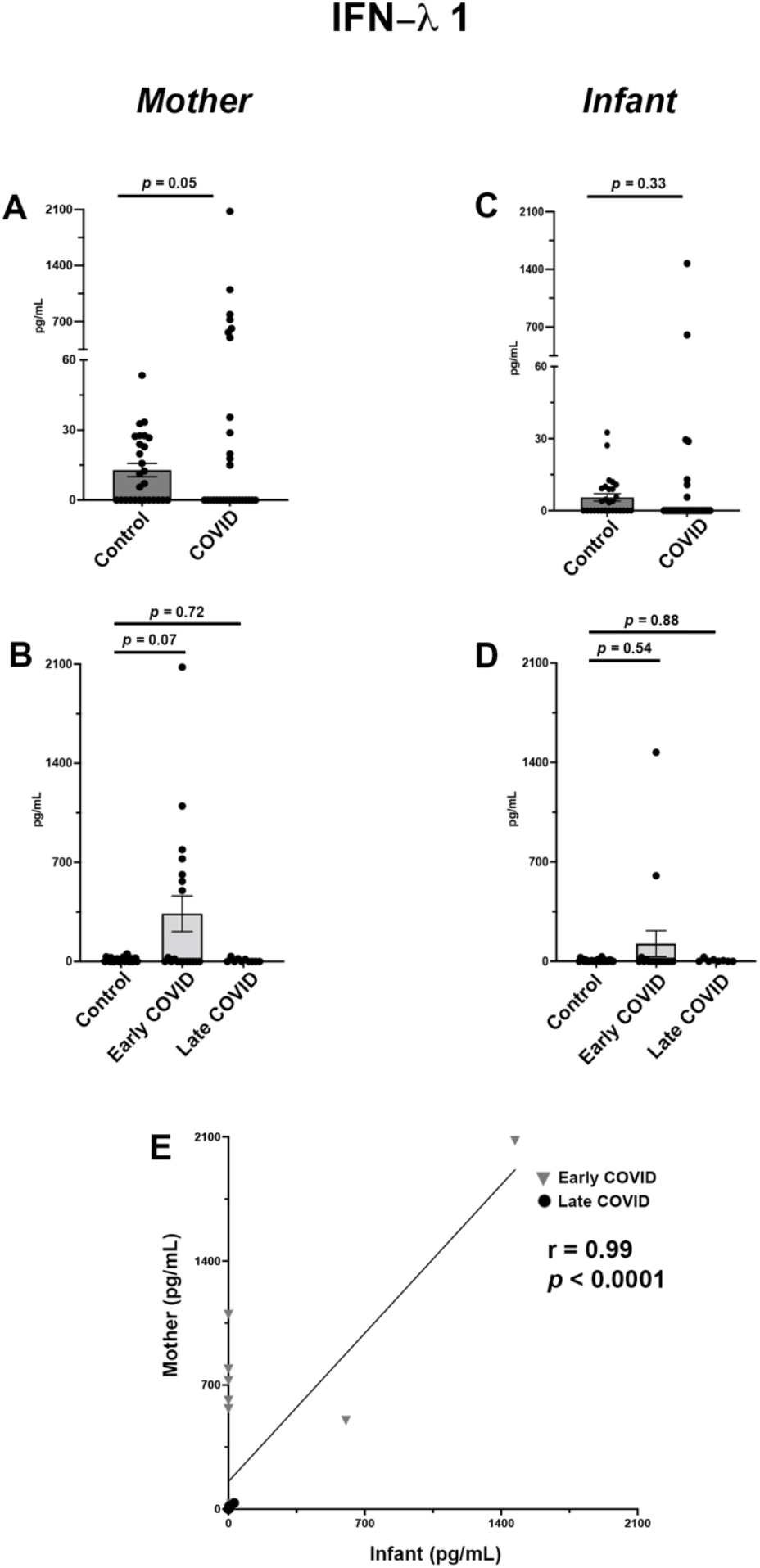
IFN-λ 1 levels in maternal and infant serum following SARS-CoV-2 infection in pregnancy. A: Maternal serum IFN-λ 1 expression in Control vs COVID cohorts. B: Maternal serum IFN-λ 1 sub-analysis among Control, Early COVID and Late COVID cohorts. C. Infant serum IFN-λ 1 expression in Control vs COVID cohorts. D: Infant serum IFN-λ 1 sub-analysis among Control, Early COVID and Late COVID cohorts. E: Correlation analysis of IFN-λ 1 expression in maternal-infant dyads. *Control*, *COVID*, *Early COVID*, and *Late COVID*: as described in Figures 1 and 2.

**Supplemental Table 1:** COVID-19 versus control maternal blood and cord blood cytokine levels.

| Cytokine | MOM BLOOD |  |  |  | CORD BLOOD / INFANT BLOOD |  |  |  |
| --- | --- | --- | --- | --- | --- | --- | --- | --- |
|  | Mean (SD) |  |  |  | Mean (SD) |  |  |  |
|  | CONTROL<br>N=28 | COVID<br>N=31 | MD<br>(95% CI) | P-Value | CONTROL<br>N=28 | COVID<br>N=31 | MD (95%<br>CI) | P-Value |
| **IL-1β** | 20.7<br>(44.0) | 144.6<br>(319.3) | 123.8<br>(-95.2,<br>342.9) | 0.23 | 69.1<br>(274.9) | 276.9<br>(853.3) | 207.8<br>(-889.0,<br>1304.6) | 0.66 |
| **IL-6** | 26.9<br>(57.9) | 544.7<br>(887.9) | 517.9<br>(125.8,<br>909.9) | **0.01** | 34.4<br>(65.5) | 610.2<br>(652.3) | 575.8<br>(293.7,<br>857.9) | **0.0002** |
| **TNF-α** | 19.8<br>(46.5) | 3009.2<br>(n/c) | n/c | n/c | 7.1<br>(14.2) | 119.6<br>(393.1) | 112.4<br>(-706.3,<br>931.2) | 0.72 |
| **IP-10** | 95.8<br>(91.5) | 598.3<br>(448.3) | 502.5<br>(329.3,<br>675.7) | **<0.0001** | 74.4<br>(60.9) | 448.2<br>(353.9) | 373.7<br>(239.7,<br>507.7) | **<0.0001** |
| **IFN-λ1** | 21.1<br>(12.4) | 271.6<br>(481.0) | 250.5<br>(-4.1,<br>505.1) | 0.05 | 10.5<br>(8.3) | 88.1<br>(293.3) | 77.7<br>(-82.2,<br>237.5) | 0.33 |
| **IL-8** | 57.8<br>(220.7) | 284.0<br>(415.9) | 226.2<br>(-37.7,<br>133.9) | 0.09 | 137.7<br>(286.2) | 2792.3<br>(5631.6) | 2654.7<br>(338.24,<br>4971.1) | **0.0256** |
| **IL-12p70** | 5.2<br>(12.1) | 38.7<br>(149.2) | 33.5<br>(-66.9,<br>113.9) | 0.47 | 3.7<br>(5.7) | 249.0<br>(23.4) | 245.3<br>(171.9,<br>318.6) | **0.005** |
| **IFN-α2** | 18.1<br>(34.2) | 140.7<br>(442.6) | 112.5<br>(-121.6,<br>66.7) | 0.31 | 2.8<br>(5.9) | 86.2<br>(368.3) | 83.4<br>(-476.7,<br>643.5) | 0.72 |
| **IFN-λ2/3** | 126.5 (159.6) | 922.6<br>(2224.7) | 796.1<br>(-780.0,<br>2372.2) | 0.30 | 117.2<br>(56.4) | 478.3<br>(1812.8) | 361.1<br>(-1857.9,<br>2580.0) | 0.70 |
| **GM-CSF** | 19.1<br>(35.2) | 89.3<br>(368.9) | 70.2<br>(-268.6,<br>409.1) | 0.65 | 12.8<br>(21.0) | n/c | n/c | n/c |
| **IFN-β** | 36.2<br>(51.8) | 253.4<br>(293.2) | 217.2<br>(9.0,<br>425.4) | **0.04** | 74.3<br>(246.4) | 130.6<br>(224.3) | 56.2<br>(-251.3,<br>363.7) | 0.70 |
| **IL-10** | 10.3<br>(24.7) | 206.6<br>(442.0) | 196.3<br>(-69.9,<br>462.6) | 0.14 | 6.4<br>(14.5) | 87.3<br>(193.8) | 80.9<br>(-66.5,<br>228.2) | 0.26 |
| **IFN-γ** | 25.2<br>(70.8) | 80.4<br>(338.4) | 55.2<br>(-222.8,<br>333.1) | 0.66 | 17.3<br>(31.8) | 8.1<br>(2.2) | -9.2<br>(-47.7, 29.2) | 0.54 |

**Abbreviations:** MD = mean difference; CI = confidence interval; IL = interleukin; TNF = tumor necrosis factor; IFN = interferon; GM-CSF = granulocyte-macrophage colony-stimulating factor; IP = interferon gamma-induced protein; n/c = not able to calculate

**Supplemental Table 2:** Maternal blood and cord blood cytokine comparisons by trimester of maternal infection.

| Cytokine | Control<br>N=28<br>Mean (SD) | 1<sup>st</sup> / 2<sup>nd</sup><br>Trimester<br>COVID<br>N=18<br>Mean (SD) | 3<sup>rd</sup> Trimester<br>COVID<br>N=10<br>Mean (SD) | P-value<br>Control<br>vs 1<sup>st</sup>/2<sup>nd</sup> | P-Value<br>Control<br>vs 3<sup>rd</sup> | P-Value<br>1<sup>st</sup>/2<sup>nd</sup> vs<br>3<sup>rd</sup> |
| --- | --- | --- | --- | --- | --- | --- |
| **Maternal Blood** |  |  |  |  |  |  |
| **IL-1β** | 20.7 (44.0) | 176.3 (390.7) | n/c | 0.67 | n/c | n/c |
| **IL-6** | 26.9 (57.9) | 667.4 (1027.1) | 328.2 (435.5) | **0.02** | 0.09 | 0.38 |
| **TNF-α** | 19.7 (46.5) | n/c | n/c | n/c | n/c | n/c |
| **IP-10** | 95.8 (91.5) | 370.1 (646.1) | 593.0 (484.2) | 0.08 | **0.009** | 0.35 |
| **IFN-λ 1** | 21.1 (12.4) | 441.2 (554.4) | 23.4 (8.7) | 0.07 | 0.72 | 0.27 |
| **IL-8** | 57.8 (220.7) | 346.8 (474.7) | 164.9 (227.4) | 0.05 | 0.31 | 0.32 |
| **IL-12p70** | 5.2 (12.1) | 349.9 (97.3) | n/c | 0.13 | n/c | n/c |
| **IFN-α2** | 18.1 (34.2) | 177.7 (520.9) | 72.0 (211.5) | 0.45 | 0.70 | 0.75 |
| **IFN-λ 2/3** | 126.5 (159.6) | 1347.6 (2583.5) | 333.0 (819.8) | 0.26 | 0.78 | 0.62 |
| **GM-CSF** | 19.1 (35.2) | 179.8 (437.3) | n/c | 0.59 | n/c | n/c |
| **IFN-β** | 36.2 (51.8) | 307.0 (337.6) | n/c | **0.02** | n/c | n/c |
| **IL-10** | 10.3 (24.7) | 447.1 (391.2) | 73.6 (214.9) | **0.04** | 0.66 | 0.18 |
| **IFN-γ** | 25.2 (70.8) | 119.1 (475.4) | n/c | 0.83 | n/c | n/c |
| **Cord Blood** |  |  |  |  |  |  |
| **IL-1β** | 69.1 (274.9) | 1253.7 (810.5) | 85.4 (6.3) | 0.29 | 0.91 | 0.18 |
| **IL-6** | 34.4 (65.5) | 634.1 (580.5) | 567.1 (776.7) | **0.0007** | 0.15 | 0.83 |
| **TNF-α** | 7.1 (14.2) | 218.5 (464.2) | n/c | 0.51 | n/c | n/c |
| **IP-10** | 74.4 (60.9) | 514.1 (334.6) | 406.3 (375.8) | **<0.0001** | **0.04** | 0.48 |
| **IFN-λ 1** | 10.5 (8.3) | 130.3 (351.7) | 11.4 (9.3) | 0.54 | 0.88 | 0.59 |
| **IL-8** | 137.7 (286.2) | 2690.3 (5471.3) | 3009.2<br>(6343.0) | 0.07 | 0.24 | 0.90 |
| **IL-12p70** | 3.7 (5.7) | 251.3 (28.7) | n/c | 0.05 | n/c | n/c |
| **IFN-α2** | 2.8 (5.9) | 129.9 (448.8) | n/c | 0.67 | n/c | n/c |
| **IFN-λ 2/3** | 117.2 (56.3) | 700.3 (2207.4) | n/c | 0.69 | n/c | n/c |
| **GM-CSF** | 12.8 (21.0) | n/c | n/c | n/c | n/c | n/c |
| **IFN-β** | 74.3 (246.4) | 171.1 (284.2) | 53.6 (102.9) | 0.56 | 0.89 | 0.60 |
| **IL-10** | 6.4 (14.5) | 114.8 (229.9) | 35.5 (65.7) | 0.35 | 0.53 | 0.50 |
| **IFN-γ** | 17.3 (31.8) | n/c | 9.8 (1.9) | n/c | 0.80 | n/c |

**Abbreviations:** MD = mean difference; CI = confidence interval; IL = interleukin; TNF = tumor necrosis factor; IFN = interferon; GM-CSF = granulocyte-macrophage colony-stimulating factor; IP = interferon gamma induced protein; n/c = not able to calculate

**Supplemental Table 3:** Correlation between maternal blood and cord blood cytokine levels.

| Cytokine | Correlation coefficient<br>COVID group only | P-Value |
| --- | --- | --- |
| **IL-1β** | n/c | n/c |
| **IL-6** | -0.04 | 0.86 |
| **TNF-α** | n/c | n/c |
| **IP-10** | 0.78 | **<0.0001** |
| **IFN-λ 1** | 0.99 | **<0.0001** |
| **IL-8** | -0.16 | 0.51 |
| **IL-12p70** | n/c | n/c |
| **IFN-α2** | 0.99 | 0.06 |
| **IFN-λ 2/3** | n/c | n/c |
| **GM-CSF** | n/c | n/c |
| **IFN-β** | 0.19 | 0.76 |
| **IL-10** | 0.71 | 0.12 |
| **IFN-γ** | n/c | n/c |

